# Circulating androgen regulation by androgen-catabolizing gut bacteria in male mouse gut

**DOI:** 10.1101/2022.07.20.500890

**Authors:** Tsun-Hsien Hsiao, Chia-Hong Chou, Yi-Lung Chen, Po-Hsiang Wang, Guo-Jie Brandon-Mong, Tzong-Huei Lee, Tien-Yu Wu, Po-Ting Li, Chen-Wei Li, Yi-Li Lai, Yu-Lin Tseng, Chao-Jen Shih, Mei-Jou Chen, Yin-Ru Chiang

## Abstract

Abnormally high circulating androgen levels have been considered a causative factor for benign prostatic hypertrophy and prostate cancer. Recent studies suggested that gut bacteria can alter sex steroid profile of host; however, the underlying mechanisms and bacterial taxa remain elusive. *Thauera* sp. strain GDN1 is an unusual betaproteobacterium capable of aerobic and anaerobic androgen catabolism in environmental conditions (37°C) resembling the mammalian gut. The strain GDN1 administration to C57BL/6J mice through oral gavage profoundly affected gut bacterial community, along with an approximately 50% reduction in serum androgen level in male mice. Our RT–qPCR results revealed the differential expression of aerobic and anaerobic androgen catabolic genes in the mouse ileum (microaerobic) and caecum (anaerobic), respectively. Furthermore, androgenic ring-cleaved metabolites were detected in the mouse fecal extract. This study discovered that androgen serves as a carbon source of gut microbes and that androgen-catabolizing gut bacteria can modulate host circulating androgen levels.

**Highlights:**

- *Thauera* sp. strain GDN1 administration through oral gavage regulated mouse serum androgen levels.
- The biochemical, genetic, and metabolite profile analyses revealed the occurrence of bacterial androgen catabolism in the mouse gut.
- Androgen catabolism proceeds through the O_2_-dependent and O_2_-independent catabolic pathways in mouse ileum and caecum, respectively.
- A possibility to harness *Thauera* sp. strain GDN1 as a functional probiotic to treat hyperandrogenism.

**Graphical Abstract:** 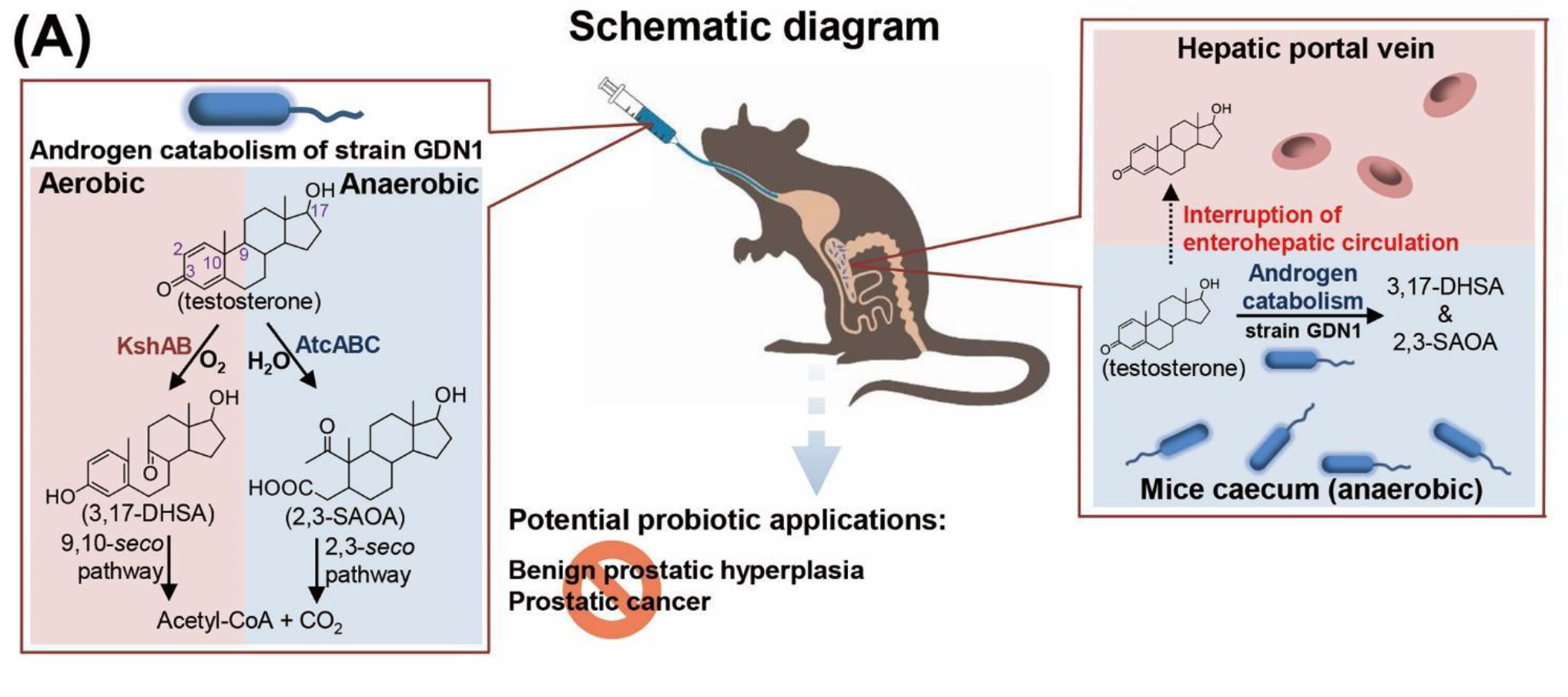

**In brief:** Hsiao et al. found that oral administration of androgen-catabolizing *Thauera* species regulated mouse serum androgen level. They characterized the gut microbe–mediated androgen catabolism through genetic and biochemical analyses. Their discovery portends a possibility of harnessing androgen-catabolic gut bacteria as functional probiotics to treat hyperandrogenism.

## Introduction

Sex steroids, such as androgens, modulate vertebrate physiology, development, reproduction, and behavior (Doyle and Meeks, 2018). Vital androgens include testosterone, dihydrotestosterone (DHT), and androstenedione. Androgens have been considered a critical mediator of androgenetic alopecia (Trüeb, 2002), benign prostatic hypertrophy (Colldén et al., 2019; Russell and Wilson, 1994) and prostate cancer (Pernigoni et al., 2021) in men. An excess of circulating androgens can cause polycystic ovary syndrome in women (Wu et al., 2017).

In mammals, sex steroids are excreted through either feces or urine; for instance, male mice excrete significantly more androgens through feces (approximately 60%) than female mice (<50%) (Auer et al., 2020). Steroids are recycled between the liver and the gut through enterohepatic circulation. Cholesterol and bile acids may be recycled more than 10 times before being excreted through feces (Adlercreutz et al., 1979; Cai and Chen, 2014); the reabsorption of these steroids occurs mainly in human small intestine (Lu et al., 2001) as well as rodent small intestine and caecum (Hugenholtz and de Vos, 2018; Nakayama et al., 1999). By contrast, little is known regarding the metabolic fate and flux of sex steroids in the mammalian gut. Current knowledge indicates that androgens undergo glucuronidation in the liver and are then delivered into the gut. The conjugated androgens become less susceptible to reabsorption and are eventually excreted through feces (Cross et al., 2018). Recent studies have also reported that gut microbes can mediate deglucuronidation, which increases free androgen levels in the mouse caecum and human colon (Colldén et al., 2019). The deconjugated androgens, mainly testosterone and DHT, are present in the bacteria-rich colon of young men at concentrations much higher than those in the serum (Colldén et al., 2019). Most gut androgens (approximately 80%) are reabsorbed into the blood through enterohepatic circulation (Li et al., 2022; Sandberg and Slaunwhite, 1956).

Sex steroids may be involved in bidirectional metabolic interactions between bacteria and their vertebrate hosts (Neuman et al., 2015; Vom Steeg and Klein, 2017); however, the underlying mechanisms and bacterial taxa remain largely unknown. Understanding these underlying mechanisms warrants the examination of host–microbe interactions, particularly the examination of the microbial metabolic contributions to the host metabolome, at the molecular level (Crowe et al., 2018; Koppel and Balskus, 2016; Pandey and Sassetti, 2008; Ridlon, 2020; Van der Geize et al., 2007). There is some evidence that sex may influence the diversity, composition, and function of gut bacterial microbiota (Ridlon, 2020), although the results are inconsistent. The contribution of the gut microbiota to the host steroid metabolome and to the host endocrine system might be the key in understanding the role of sex as a biological variable under both healthy and disease conditions. For instance, a recent gut microbiome study revealed that some gut microbes (e.g., *Ruminococcus gnavus*) contribute to endocrine resistance in castration-resistant prostate cancer through the transformation of pregnenolone (precursor of steroid hormones) to androgens (Pernigoni et al., 2021), increasing circulating androgen levels through enterohepatic circulation and thus enhancing prostate cancer growth in the host.

The bacterial catabolism of sex steroids has been characterized at the molecular level [see (Chiang et al., 2020) for a recent review]. Aerobic androgen catabolism through the steroid 9,10-*seco* pathway has been well characterized (Bergstrand et al., 2016; Holert et al., 2018; Horinouchi et al., 2018); it includes 3,17-dihydroxy-9,10-*seco*-androsta-1,3,5(10)-triene-9-one (3,17-DHSA) (Dresen et al., 2010) and the 3-ketosteroid 9α-hydroxylase gene *kshAB* (Capyk et al., 2011) as characteristic androgen metabolite and degradation genes, respectively. By using several denitrifying proteobacteria as the model organisms, we previously established an anaerobic degradation pathway (the steroid 2,3-*seco* pathway) for androgens; moreover, the 17β-hydroxy-1-oxo-2,3-*seco*-androstan-3-oic acid (abbreviated as 2,3-SAOA) (Wang et al., 2013) and *atcABC* encoding the 1-testosterone hydratease/dehydrogenase (Yang et al., 2016) were identified as the characteristic androgen metabolite and degradation genes for this anaerobic pathway, respectively. However, androgen catabolism has not been reported in the animal gut. Facultative anaerobes *Thauera* spp. are widely distributed in O_2_-limited environments, such as in the estuarine sediment (Shih et al., 2017) and denitrifying sludge (Yang et al., 2016) as well as in chicken (Zhang et al., 2018), pig (Zhang et al., 2018), duck (Li et al., 2021), and human (Cussotto et al., 2019; Maini Rekdal et al., 2020; Xu et al., 2017) intestines. Thus far, at least two *Thauera* species have been identified as anaerobic androgen degraders (Shih et al., 2017; Yang et al., 2016).

In this study, we used *Thauera* sp. strain GDN1 (Shih et al., 2017) (referred as strain GDN1 hereafter) as the model microorganism because (i) strain GDN1 can utilize major androgens as the sole carbon sources and as electron donors under both aerobic and anaerobic (denitrifying) conditions, (ii) strain GDN1 can efficiently degrade testosterone in environmental conditions (37°C and pH 6.0) resembling the mammalian gut, and (iii) *Thauera* spp. are often detected in the animal gut, but their functions in their hosts remain unknown. We performed tiered functional genomic analyses to identify the characteristic metabolites and degradation genes of strain GDN1 involved in aerobic and anaerobic androgen catabolism. Subsequently, we administered mice with strain GDN1 through oral gavage and monitored its colonization and androgen-catabolic activities in mouse intestines (**Fig. 1*A***). We also collected fecal samples from mice with different treatments and examined how the gut bacterial communities of mice varied across different treatments. Finally, the changes in host circulating androgen levels and profiles were determined using enzyme-linked immunosorbent assay kit (ELISA) and ultraperformance liquid chromatography–high-resolution mass spectrometry (UPLC–HRMS), respectively.

**Figure 1.**
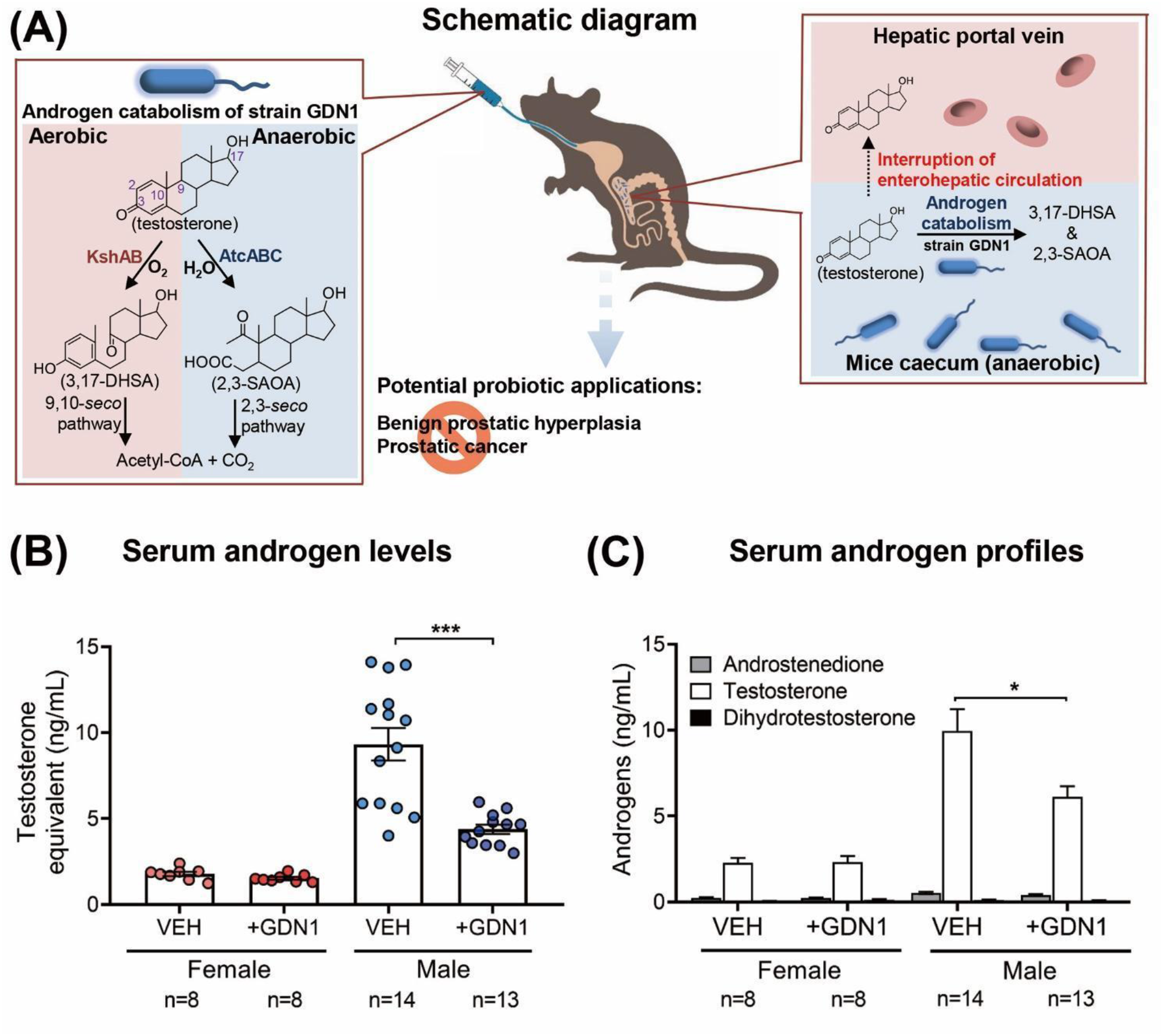
Administration of androgen-catabolic *Thauera* sp. strain GDN1 into mouse gut through oral gavage can regulate host circulating androgen levels. (**A**) Schematic of the androgen-mediated *Thauera*–mouse interactions. Strain GDN1 can catabolize androgens such as testosterone aerobically and anaerobically, and both the aerobic and anaerobic ring-cleaved metabolites (3,17-DHSA and 2,3-SAOA, respectively) were detected in the mouse gut. (**B**) Administration of male mice with strain GDN1 through oral gavage for 25 days reduces serum testosterone levels considerably. (C) Strain GDN1 administration does not affect the serum androgen metabolite profiles of male and female mice considerably. Testosterone is the most dominant serum androgen (> 90%, w/w) in all the included mice. Results are representative of three individual experiments. Statistical results were calculated with unpaired nonparametric *t*-test; \**p* < 0.05, \*\*\**p* < 0.001. All data are shown as means ± SEM of 8–14 mouse individuals.

## Results

### Physiological and multi-omics analyses for strain GDN1 growth and androgen catabolism

The mammalian gut is an O_2_-limited environment, but often with a low abundance of facultative anaerobic and microaerobic microorganisms (Ley et al., 2008). Thus far, only four bacterial strains, namely *Steroidobacter denitrificans* DSMZ18526 (Leu et al., 2011), *Sterolibacterium denitrificans* DSMZ13999 (Wang et al., 2014), *Thauera terpenica* 58Eu (Yang et al., 2016), and strain GDN1 (Shih et al., 2017), are reportedly able to anaerobically catabolize testosterone at 25°C–28°C and neutral pH (6.5–7.5). First, we characterized the physiology of these facultative anaerobes and evaluated their colonization and androgen-catabolic potentials in the mammalian gut. We anaerobically grew these bacteria with testosterone at 37°C and pH 6.0, and only strain GDN1 apparently degraded testosterone within 2 days (**Fig. 2*A***). We observed that substrate consumption accompanied a decrease in androgenic activity as well as nitrate reduction with time. Moreover, we detected temporal production of the androgenic metabolite 2,3-SAOA in the denitrifying bacterial culture (**Fig. 2*B***; left panel). However, strain GDN1 could not degrade testosterone anaerobically in the chemically defined medium without exogenous nitrate (**Fig. 2*B***; right panel), indicating the incapability of androgen catabolism under fermentative conditions. We also observed aerobic testosterone catabolism and decreased androgenic activity in a phosphate-buffered minimal medium, with 3,17-DHSA being the characteristic androgen metabolite (**Fig. 2*C***). These ring-cleaved metabolites are specific for androgen catabolism (Dresen et al., 2010; Wang et al., 2013) and exhibit negligible androgenic activity (Wang et al., 2014). These androgenic metabolites are thus suitable molecular markers for monitoring bacterial androgen catabolism in the mammalian gut. In addition to testosterone, strain GDN1 was able to grow with other major androgens including DHT and androstadienedione (1 mM for each) in a denitrifying minimal medium at 37°C and pH 6.0 (**Fig. 2*D***; left panel). By contrast, strain GDN1 could not degrade other sex steroids including progesterone and 17β-estradiol (**Fig. 2*D***; right panel). The physiological test results thus support strain GDN1 growth in the mammalian gut and suggest its metabolic specificity for androgen metabolism.

**Figure 2.**
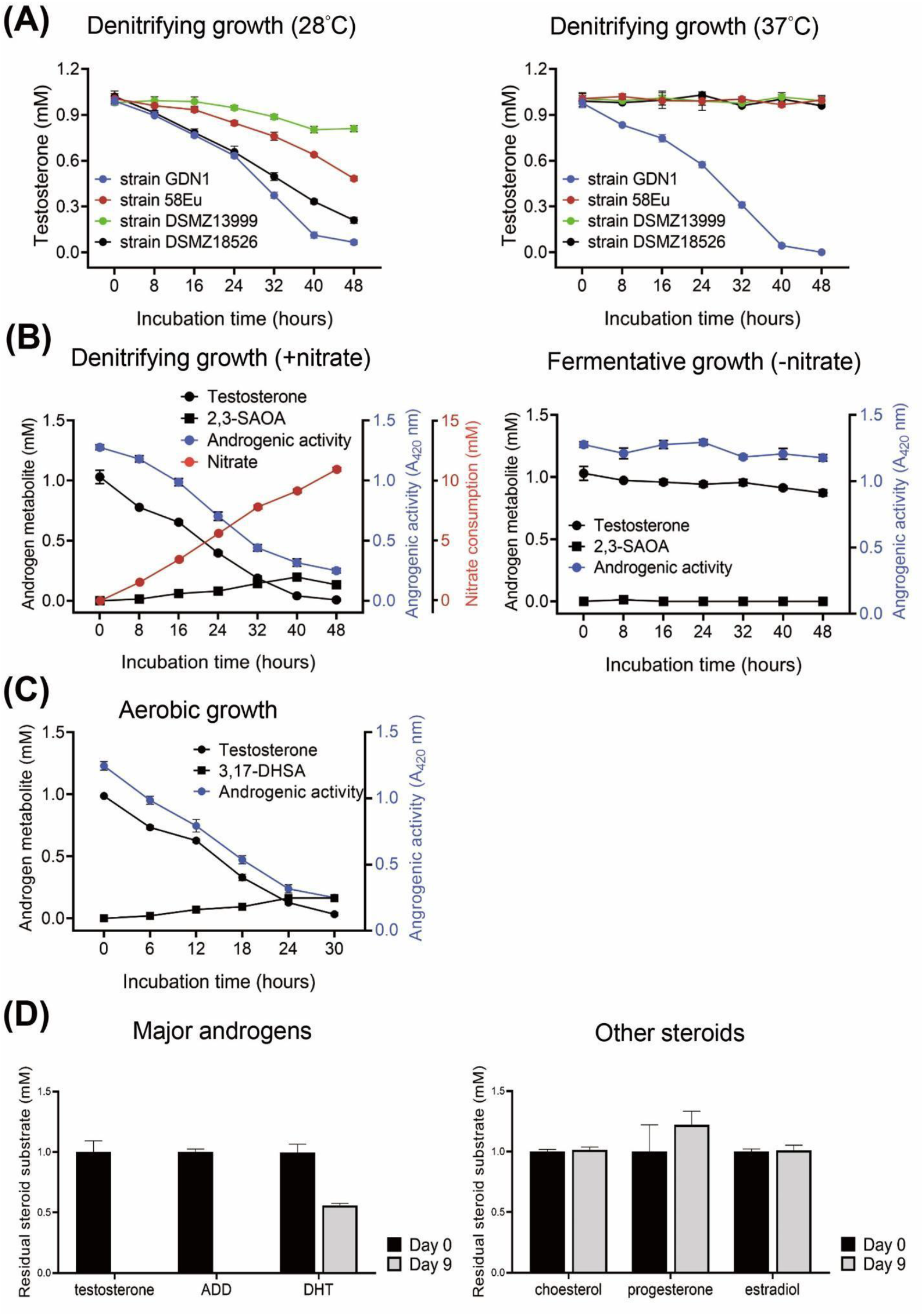
Physiology and steroid utilization of strain GDN1. (**A**) Denitrifying growth of several androgen-catabolic anaerobes with testosterone at 28°C and 37°C. (**B**) Nitrate is required for anaerobic androgen catabolism of strain GDN1. Androgenic activity in bacterial cultures was determined using a yeast androgen bioassay. (**C**) Aerobic androgen catabolism of strain GDN1. (**D**) Steroid utilization patterns of strain GDN1. Androgen metabolites were detected and quantified through UPLC–ESI–HRMS. Results are representative of three individual experiments. All data are shown as mean ± SEM of three technical replicates. Abbreviations: androstadienedione, ADD; dihydrotestosterone, DHT.

We then elucidated the biochemical mechanisms underlying androgen catabolism in strain GDN1 and identified the potential biomarkers of androgen catabolism through genomic approaches. We could identify gene clusters specific for aerobic and anaerobic androgen catabolism (**Fig. 3*A***; see detailed genomic analysis in **SI Results**). Of these, the gene products (namely the bifunctional 1-testosterone hydratase/dehydrogenase) of *atcABC* (CKCBHOJB_02409–2411; **Dataset S1**) mediate O_2_-independent androgenic A-ring activation by adding a hydroxyl group at C-1 of testosterone (Yang et al., 2016), with 2,3-SAOA as the downstream product. By contrast, the 3-ketosteroid 9α-hydroxylase gene (*kshAB*; CKCBHOJB_02264 and 02279; **Dataset S1**) is responsible for the O_2_-dependent cleavage of the androgen B-ring (Capyk et al., 2011), with 3,17-DHSA as the product.

**Figure 3.**
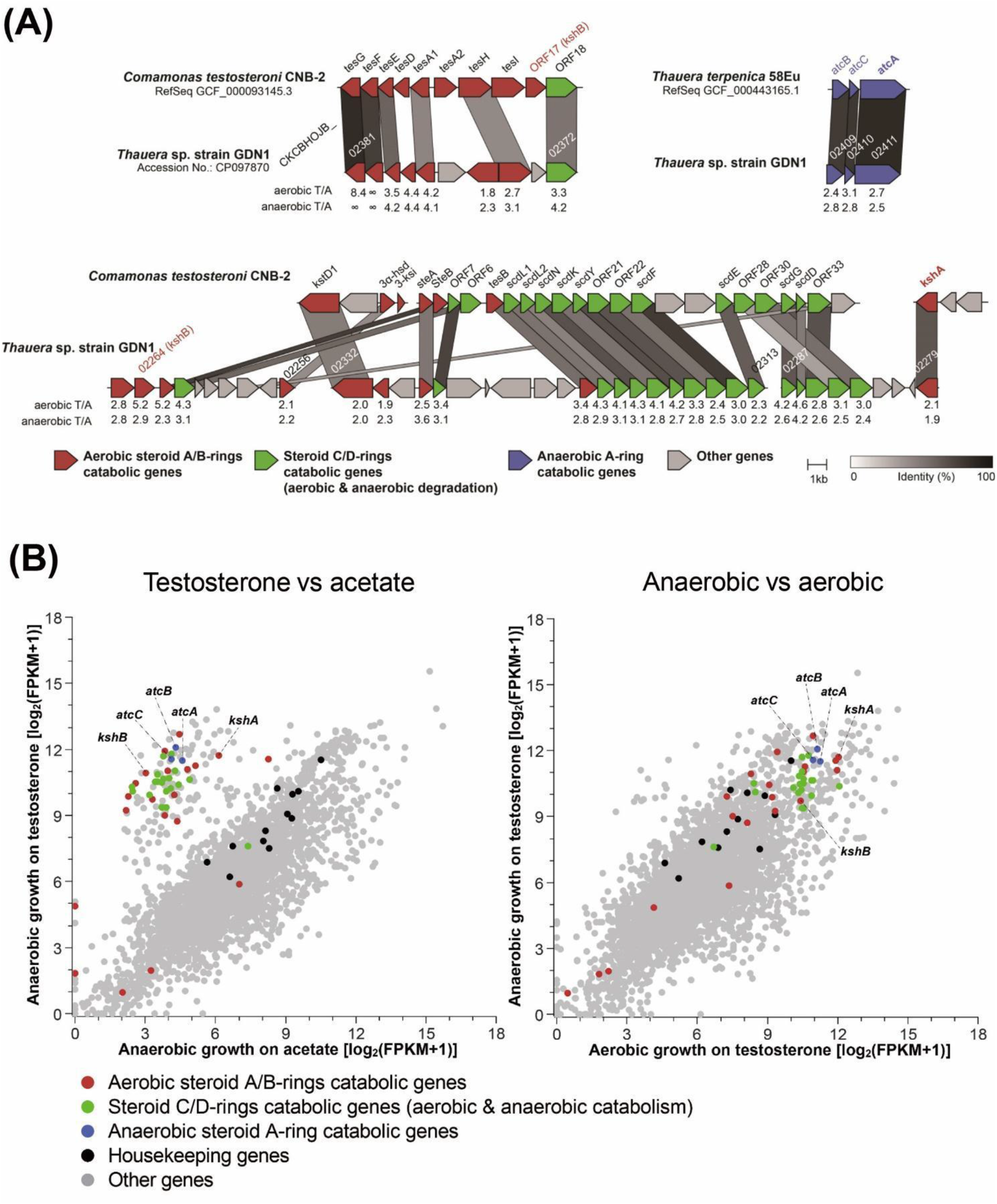
Identification and expression of androgen catabolism genes of strain GDN1. (A) Gene synteny of androgen catabolism in the strain GDN1 chromosome (accession No.: CP 097870). Strain GDN1 uses the gene products of *atcABC* and *kshAB* to catabolize androgen anaerobically and aerobically, respectively. ORFs homologous between different bacterial strains are connected by columns with different gray scaling, according to the similarities of deduced amino acid sequences. Numbers below individual ORFs of strain GDN1 indicate the gene expression ratios, which are derived from the logarithmic transformation of differential gene expression levels of strain GDN1 grown with acetate (abbreviation: A) or testosterone (abbreviation: T) under aerobic and anaerobic (denitrifying) conditions. Androgen catabolism genes are named according to those characterized in two androgen-degrading bacteria: *Comamonas testosteroni* strain CNB-2 (aerobic androgen degrader) and *Thauera terpenica* 58Eu (anaerobic androgen degrader). Synteny analysis was conducted using Clinker v0.023 (https://github.com/gamcil/clinker). (**B**) Global gene expression profiles (RNA-Seq) of strain GDN1 grown under different conditions [anaerobic growth with testosterone or acetate (left panel); anaerobic or aerobic growth with testosterone (right panel)]. Each spot represents a gene. Gene expression levels were estimated using fragments per kilobase of transcript per million mapped reads (FPKM).

To identify the inducers of these androgen catabolism genes, we incubated strain GDN1 with acetate (10 mM) or testosterone (2 mM) under aerobic and anaerobic conditions. The transcriptomes of strain GDN1 cells were then sequenced on an Illumina-based platform. We detected the distinguishable expression of *atcA* and *kshA* in the testosterone-grown cells, regardless of the oxygen conditions [**Figs. 3*B*** (right panel); see detailed transcriptome analysis in **SI Results**]. However, we observed low expression of these androgen catabolism genes in the acetate-grown strain GDN1 cells (**Figs. 3*B***; left panel). The expression levels of *atcA* and *kshA* in the same conditions were also confirmed using reverse transcription (RT)-quantitative polymerase chain reaction (qPCR) analysis (**Fig. S1**). Our multi-omics data thus indicated that (i) strain GDN1 can aerobically and anaerobically catabolize androgen through the established 9,10-*seco* (Yam et al., 2009) and 2,3-*seco* (Wang et al., 2013) pathways, respectively, and (ii) the gene expression of the androgen catabolism genes *atcABC* (for anaerobic catabolism) and *kshAB* (for aerobic catabolism) is induced by the substrate (androgen). The subunits AtcA and KshA contain molybdopterin (Yang et al., 2016) and a mononuclear iron center (Capyk et al., 2009), respectively—which are essential cofactors for their catalytic activities. Therefore, we believe that *atcA* and *kshA* are suitable biomarkers for monitoring the anaerobic and aerobic androgen catabolism, respectively, in the animal gut.

### Administration of strain GDN1 to mice through oral gavage and resulting changes in host serum androgen levels

Strain GDN1, a facultative anaerobic bacterium, was aerobically grown in tryptone soy broth, and the cells were harvested in the log phase [optical density at 600 nm (OD_600_) = 0.5–0.7]. These cells were resuspended in a basal mineral medium and stored at 4°C (within 5 days) before use. The resting strain GDN1 suspension (200 μL; containing approximately 5 × 10^8^ colony-forming units) was administered into mouse gut through oral gavage twice per week. C57BL/6J mice (aged 7 weeks) were used as the model host. All the included mice were treated with exogenous nitrate (an electron acceptor for anaerobic androgen catabolism of strain GDN1; supplemented in drinking water at a final concentration of 2 mM). The mice were treated as follows: (i) female mice administered with nitrate (2 mM in drinking water) and with (strain GDN1 treatment; n = 8) or without (vehicle treatment; n = 8) strain GDN1 administration and (ii) male mice administered with nitrate and with (strain GDN1 treatment; n = 13) or without (vehicle treatment; n = 14) strain GDN1 administration (**Fig. 4*A***). During the continual strain GDN1 administration period, we did not observe considerable weight loss or appetite loss among our mice.

**Figure 4.**
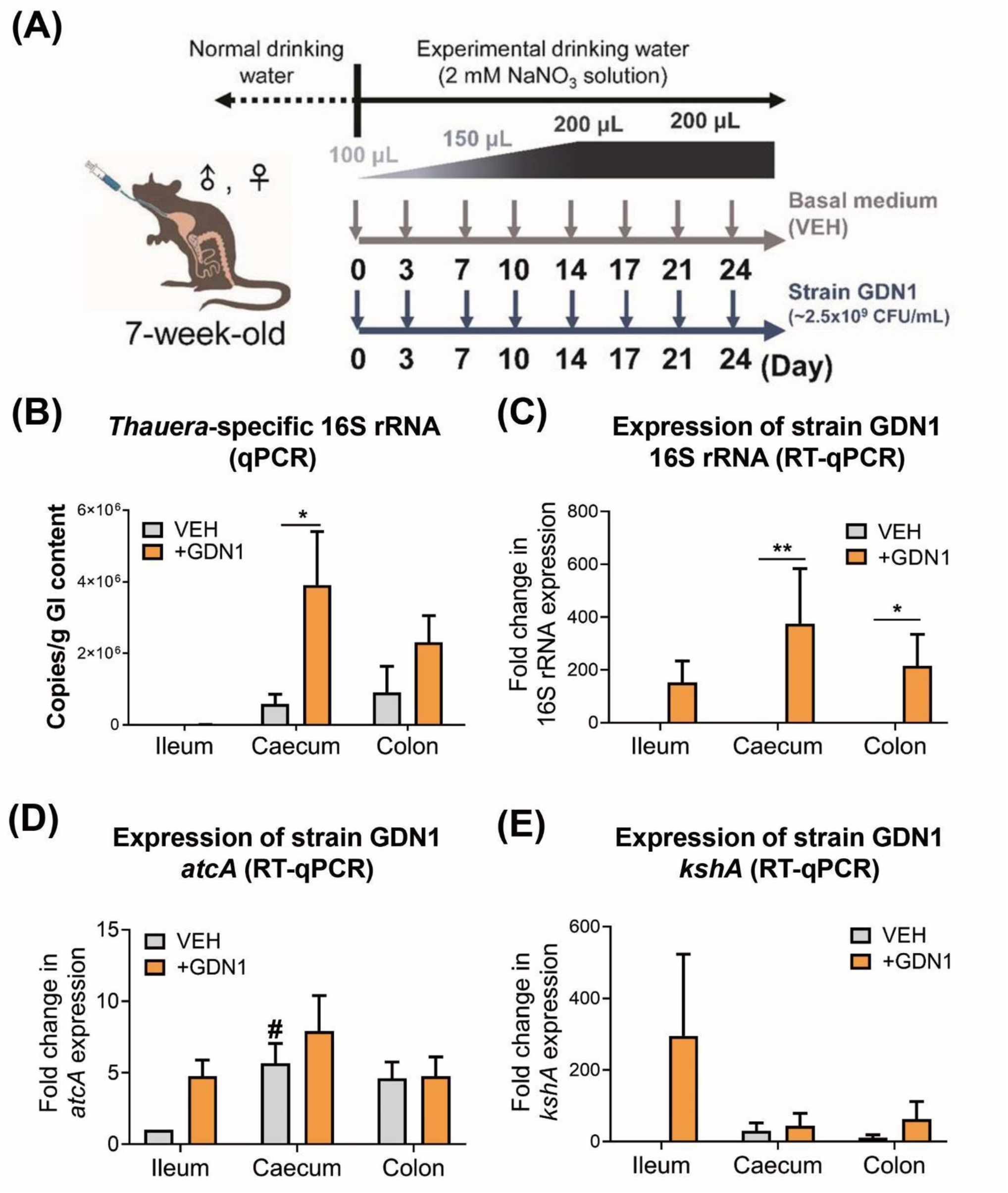
**(A)** Working flow of the administration of mice with strain GDN1 through oral gavage. Nitrate (2 mM) was supplemented in the drinking water during the period of continuous administration. (**B**) Determination of *Thauera* 16S rRNA copy number in the mouse GI contents using qPCR. All data are shown as means ± SEM of 5 randomly selected male mice. Statistical results were computed with unpaired nonparametric *t* test; \**p* < 0.05. (**C–E**) Determination of relative expression of strain GDN1-specific 16S rRNA (**C**), *atcA* (**D**), and *kshA* (**E**) in the mouse GI contents using RT-qPCR. Relative change in gene expression was calculated using the 2^−ΔΔCt^ method with the Ct value of universal 16S rRNA of bacteria as the internal control. The expression of individual genes in the vehicle-administered mouse ileum was set as 1. Results are representative of three individual experiments. Statistical results were calculated with unpaired nonparametric *t*-test; \**p* < 0.05, \*\**p* < 0.01. All the data are shown as mean ± SEM of 5 randomly selected male mice. #, expression of inherent *atcA*-like gene (see **Appendix S4** for nucleotide sequence).

We then used ELISA and UPLC–HRMS to determine the androgen levels and profiles in the mouse sera. After 25 days of continual administration of strain GDN1, we observed a considerable decrease in serum androgen levels (expressed as testosterone equivalents; see detailed information in the **Materials and Methods**) in the strain GDN1-administered male mice (**Fig. 1*B***). The ELISA results indicated that serum androgen levels (testosterone equivalents) were considerably lower in the strain GDN1-administered male mice (4.38 ± 0.27 ng/mL) than in the vehicle-administered mice (9.33 ± 0.95 ng/mL). Notably, serum androgen levels in our healthy male mice (in the vehicle group) demonstrated extreme deviation (in the range of 4–15 ng/mL), and the administration of strain GDN1 to the male mouse gut considerably reduced this deviation (in the range of 3–6 ng/mL). By contrast, we did not observe a considerable decrease in serum androgen levels in the strain GDN1-administered female mice (**Fig. 1*B***). Our data thus indicate that androgen-mediated strain GDN1– mouse interactions are gender-dependent. The serum androgen profile analysis through UPLC– HRMS indicated that testosterone (9.96 ± 1.27 ng/mL), androstenedione (0.53 ± 0.07 ng/mL), and DHT (0.12 ± 0.02 ng/mL) were the major androgens in the male vehicle-administered mice. Of these, only testosterone (6.11 ± 0.63 ng/mL after strain GDN1 treatment) considerably decreased in the strain GDN1-administered male mice (**Fig. 1*C***). By contrast, we did not observe apparent differences in the serum androgen profiles between the female mice administered with vehicle (2.28 ± 0.29 ng/mL testosterone in the serum) and strain GDN1 (2.31 ± 0.36 ng/mL testosterone in the serum). In contrast to serum testosterone levels, serum estradiol levels did not significantly change after strain GDN1 administration compared with that after vehicle administration for both the male and female mice (**Fig. S2**).

### Strain GDN1 colonization and androgen catabolism gene expression in mouse gut

The changes in the serum androgen levels of the strain GDN1-administered male mice suggested that strain GDN1 can regulate circulating androgen levels in male mice. To elucidate the mechanisms underlying the gut bacteria–host interactions, we determined the colonization and androgen-catabolic activity of strain GDN1 in the mouse gut. After 25 days of administration with strain GDN1, we sacrificed the mice and extracted the bacterial DNA from the mouse gastrointestinal (GI) contents to determine the abundance of *Thauera* spp. in the ileum, caecum, and colon through qPCR. The results of qPCR using *Thauera*-specific primers suggested that *Thauera* spp. mainly colonized in the caecum [(3.9 ± 1.5) × 10^6^ copies/g of GI content] of the strain GDN1-administered male mice (**Fig. 4*B***). By contrast, *Thauera* spp. mainly colonized in the colon [(0.9 ± 0.7) × 10^6^ copies/g of GI content] of the vehicle-administered mice. Moreover, we extracted RNA from the mouse GI contents of the ileum, caecum, and colon and then performed RT-qPCR to examine the relative expression of strain GDN1-specific 16S rRNA and androgen catabolism genes by using the 2^−ΔΔCt^ method (Livak and Schmittgen, 2001). Among all vehicle- and GDN1-administered mouse GI contents, the expression of strain GDN1 16S rRNA showed the most significant activity in the caecum of the GDN1-administered male mice (375-fold higher than that in the vehicle-administered male mouse ileum; **Fig. 4*C***). By contrast, strain GDN1-specific 16S rRNA was not considerably expressed in the vehicle-administered mouse gut. These data indicated that the colonization and growth of strain GDN1 mainly occur in the mouse caecum. To confirm the viability of strain GDN1 in the mouse gut, we incubated fecal samples from the strain GDN1- and vehicle-administered mice (25 days after the first administration) with testosterone (1 mM) under denitrifying conditions. After 3 days of anaerobic incubation at 37°C and pH 6.0, apparent testosterone consumption was observed only in the denitrifying culture spiked with feces from the strain GDN1-administered mice (**Fig. 5**). These results indicate that strain GDN1 has a high survival rate in the mouse gut.

**Figure 5.**
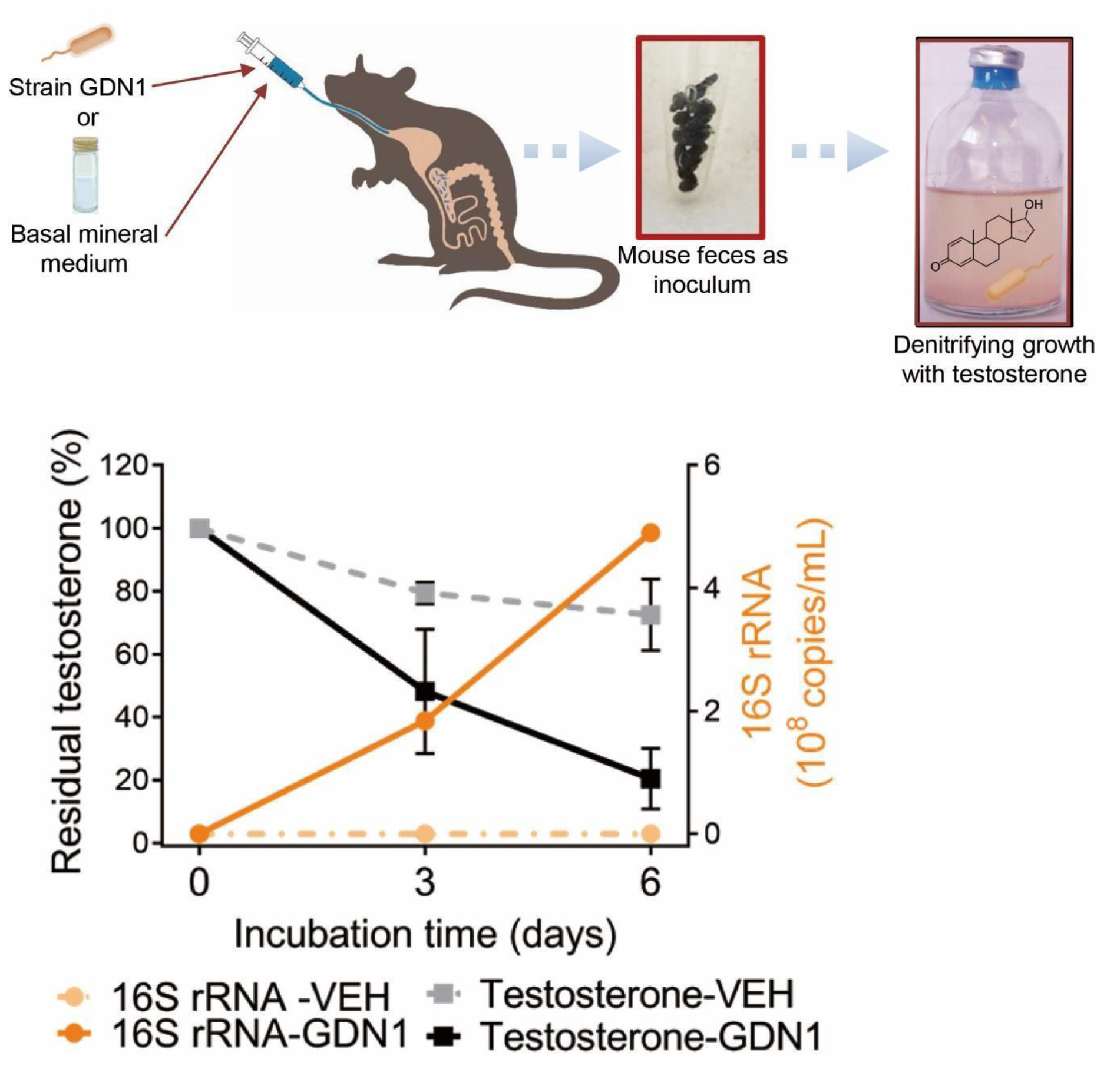
Anaerobic growth of fecal bacteria from the strain GDN1-administered mice (abbreviation: GDN1) or vehicle-administered mice (VEH) mice with testosterone (1 mM; set as 100%) as sole carbon source. After 25 days of continual administration, the fresh mouse feces (approximately 0.1 g) were collected and immediately incubation with testosterone in a denitrifying medium, and the temporal changes in substrate consumption and the copy number of *Thauera* 16S rRNA in bacterial cultures was determined. The residual testosterone concentration was examined using HPLC. The copy number of *Thauera* 16S rRNA was determined through qPCR.

We then applied RT-qPCR to investigate the expression of the androgen catabolism genes in the mouse gut. We determined *atcA* expression in the mouse gut, with the highest *atcA* expression in the caecum of the strain GDN1-administered male mice (7.92-fold higher than that in vehicle-administered male mouse ileum; **Fig. 4*D***). Notably, we also detected the expression of *kshA* in the mouse gut; *kshA* expression in the ileum of the strain GDN1-administered male mice was 294-fold higher than that in the ileum of the vehicle-administered mice (**Fig. 4*E***).

### Impact of strain GDN1 administration on mouse gut microbiota

We next sought to examine how the gut bacterial communities varied across different mouse treatments. After normalizing all samples to 48558 reads, 993457 amplicon sequence variants (ASVs) were generated, which belong to 280 bacterial genera, 118 families, and 12 phyla. In the mice administered with stain GDN1, we observed an increase in the *Thauera* population (**Fig. S3**). After 3 weeks of the first administration, the relative abundance of *Thauera* reached approximately 0.1% in the gut of both male and female mice. By contrast, the relative abundance of *Thauera* remained low (< 0.01%) in the gut of the vehicle-administrated mice.

The non-metric multidimensional scaling (NMDS) ordination indicated that the gut bacterial communities of male mice did not cluster by the strain GDN1 administration (**Fig. 6*A***; left panel), although permutational multivariate analysis of variance (PERMANOVA) based on Bray-Curtis dissimilarity matrix indicated a difference in the clustering (F-value = 2.451, global R^2^ = 0.100, *p* value = 0.026). In contrast, among female mice, the gut bacterial community structure of the strain GDN1-administered female mice (GF) clustered distinctly from that of the vehicle-administered female mice (VF) as confirmed by PERMANOVA (F-value = 2.451, global R^2^ = 0.100, *p* value = 0.026). Our data thus indicated a discernible change in the gut bacterial community structure of female mice by the strain GDN1 administration.

**Figure 6.**
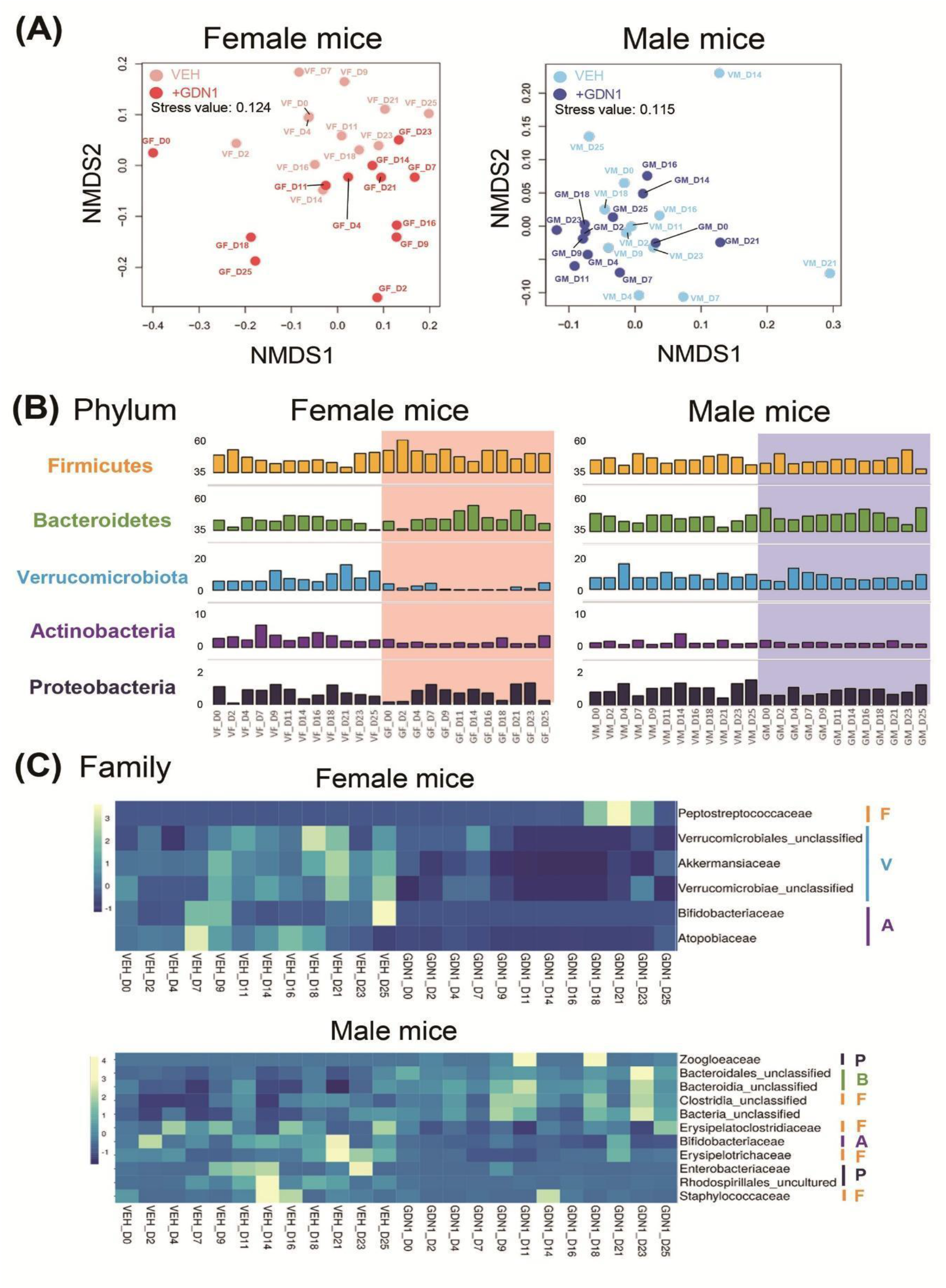
Impact of strain GDN1 administration on mouse gut microbiota. (**A**) Gut bacterial communities across the mouse treatments. NMDS analysis based on Bray-Curtis distance matrix data (genus level) was used to determine the similarities between the gut bacterial communities from different mouse treatments, including the vehicle-administered male mice (VM), strain GDN1-administered male mice (GM), vehicle-administered female mice (VF), and strain GDN1-administered female mice (GF). Numbers shown are the days after the first oral administration. (**B**) Relative abundances of the most prevalent bacterial phyla (bar length) in the mouse fecal samples. (**C**) Relative abundance changes of major bacteria families in the fecal samples from different mouse treatments. Bacterial phyla: ochre, Firmicutes (F); green, Bacteroidetes (B); purple, Actinobacteria (A); light blue, Verrucomicrobiota (V); Dark blue, Proteobacteria (P).

Of the 12 identified bacterial phyla, 5 phyla dominated the mouse gut microbiota (average cumulative abundance = 98.5%), namely Firmicutes (synonym Bacillota), Bacteroidetes (synonym Bacteroidota), Verrucomicrobiota, Actinobacteria (synonym Actinobacteriota), and Proteobacteria (synonym Pseudomonadota) (**Fig. 6*B***). Among them, Firmicutes (relative abundance = 46.7%) and Bacteroidetes (relative abundance = 42.7%) comprised the most dominant phyla in the mouse gut microbiota; Proteobacteria was the least dominant (relative abundance = 8.5%). Although all test mice were treated with 2 mM nitrate (supplemented through drinking water), we did not observe an apparent increase in the abundance of denitrifying proteobacteria. We observed an apparent decrease in the abundance of Verrucomicrobiota and Actinobacteria in the gut of strain GDN1-administered female (GF) mice (**Fig. 6*B***; left panel). Furthermore, in the gut of female mice, we identified 3 bacterial families within Verrucomicrobiota (e.g., Akkermansiaceae and 2 unclassified families) that are most affected by the strain GDN1 administration (**Fig. 6*C***; upper panel). In addition, we observed an apparent decrease in the relative abundance of the family Bifidobacteriaceae (Phylum Actinobacteria) in the GM and GF mice (**Fig. 6*C***). In the strain GDN1-administered male mice (GM), we also observed a decrease in the abundance of other families, including Erysipelatoclostridiaceae (Firmicutes), Erysipelotrichaceae (Firmicutes), Staphylococcaceae (Firmicutes), Enterobacteriaceae (Proteobacteria), and Rhodospirillales_uncultured (Proteobacteria). In the GM treatment, the strain GDN1 administration resulted in the increase in the abundance of gut bacteria belonging to Zoogloeaceae (Proteobacteria), Bacteroidales_unclassifed (Bacteroidetes), Bacteroidia_unclassified (Bacteroidetes), and Clostridia_unclassified (Firmicutes) (**Fig. 6C**; bottom panel).

### Metabolite profile analysis of fecal extracts for strain GDN1-mediated androgen catabolism in mouse gut

To monitor the microbial activity of androgen catabolism in the mouse gut, fecal samples were extracted using ethyl acetate, and the androgenic metabolites were identified through UPLC– electrospray ionization (ESI)–HRMS (**Fig. 7**). We employed extracted ion current (EIC) for *m/z* 305.21 (the most dominant ion peak of 2,3-SAOA) to detect 2,3-SAOA in the fecal extracts of our male mice. In the fecal extract of the strain GDN1-administered male mice, we identified a compound with behaviors in UPLC (retention time = 7.1 min) and ESI–MS spectrum (quasimolecular adducts [M + H]^+^ and [M + Na]^+^ at *m/z* 323.22 and 345.20, respectively) matching with those of the authentic standard 2,3-SAOA (**Fig. 7A**). By contrast, we did not detect any signals corresponding to 2,3-SAOA in the fecal extract of the vehicle-administered mice. Subsequently, we employed EIC for *m/z* 303.19 (the most dominant ion peak of 3,17-DHSA) to detect 3,17-DHSA in fecal extracts of our male mice. We detected this aerobic ring-cleaved metabolite in the fecal extract of the strain GDN1-administered male mice, with the UPLC–ESI–HRMS behaviors matching with those of the authentic standard 3,17-DHSA (**Fig. 7B**). The largest amount of 3,17-DHSA (980 ± 120 pg/g of feces) and 2,3-SAOA (70 ± 20 pg/g of feces) was determined in the mouse feces sampled after 2 weeks of the first oral administration. The metabolite profile analysis of the mouse fecal extracts thus suggests the production of both 3,17-DHSA and 2,3-SAOA (the aerobic and anaerobic ring-cleaved metabolites, respectively) in our male mice administered with strain GDN1.

**Figure 7.**
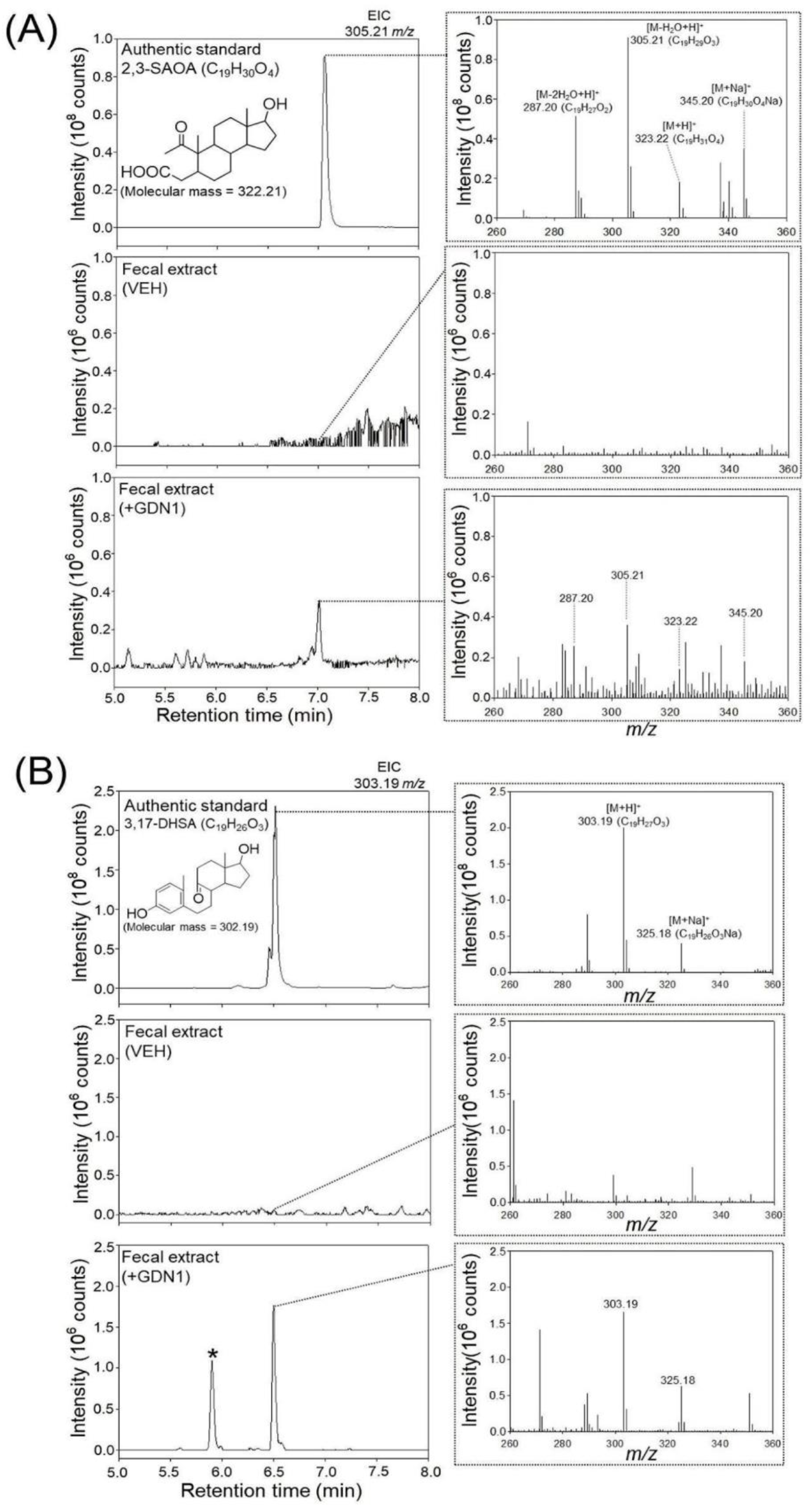
UPLC–ESI–HRMS detection of (A) 2,3-SAOA (anaerobic ring-cleaved metabolite) and (B) 3,17-DHSA (aerobic ring-cleaved metabolite) in the fecal extract of male mice administered with strain GDN1. Aforementioned ring-cleaved metabolites were not detected in the vehicle-administered mice. Fresh mouse feces were sampled 2 weeks after the first oral administration and was stored at −80°C before use. The fecal samples were extracted using ethyl acetate, and the androgen metabolites were analyzed through UPLC–ESI–HRMS. The predicted elemental composition of individual metabolite ions was calculated using MassLynx Mass Spectrometry Software (Waters); *unidentified metabolite.

## Discussion

In this study, we observed that androgen catabolism occurs in the animal gut. Surprisingly, our qPCR and PCR-based functional assay results suggested the presence of inherent *Thauera* spp. and *atcA*-like genes in the mouse gut. *Thauera* spp. are metabolically versatile and can grow with various hydrocarbons, including sugars, aromatics, and terpenoids, under aerobic, anaerobic, and fermentative conditions (Anders et al., 1995; Foss and Harder, 1998; Shih et al., 2017); these compounds are abundant in the gut of animals (Cussotto et al., 2019; Li et al., 2021; Maini Rekdal et al., 2020; Xu et al., 2017; Zhang et al., 2018). However, the function of *Thauera* spp. in the animal gut remains unclear. Among the tested sex steroids, we found that strain GDN1 can only utilize androgens. The high substrate specificity of strain GDN1 thus excludes its potential adverse effects on estrogen and progesterone metabolism in the host.

Although the expression of androgen catabolism genes of strain GDN1 are induced by testosterone, their expression at a low level was observed in the acetate-grown cells. Moreover, the androgen catabolism genes are not regulated by oxygen availability. Our bacterial cultivation data thus suggest the expression of androgenic catabolism genes in the mouse gut, regardless of substrate and oxygen availability. Indeed, we detected strain GDN1 *kshA* and *atcA* expression in the mouse gut, mainly in the ileum and caecum, respectively. The low *atcA* expression in the mouse gut was partially due to the extremely low testosterone content (< 20 ng/g feces) in the lower GI tract (Colldén et al., 2019). Moreover, we sampled bacterial RNA from the mouse GI after 25 days of continual strain GDN1 administration, which is when gut androgens may be exhausted.

The monitoring of microbial functional gene expression in the animal gut is challenging because of the highly dynamic gene expression of the gut microbiota (Booijink et al., 2010; Miro-Blanch and Yanes, 2019). As an alternative approach, we applied UPLC–HRMS to detect *KshA* and *AtcA* products in the GI contents and feces of mice. However, the levels of androgen metabolites in the GI contents of our mice were too low to detect. Nonetheless, we detected the ring-cleaved metabolites 3,17-DHSA and 2,3-SAOA in the fecal extracts of the strain GDN1-administered mice, suggesting that both aerobic and anaerobic androgen catabolism occur in the mouse gut. Given that oxygenase-mediated steroid metabolism can proceed under microaerobic conditions (Waldbauer et al., 2011), aerobic androgen catabolism might play a crucial role in gut androgen consumption, particularly in the ileum. Therefore, we believed that *kshA*-mediated aerobic androgen catabolism mainly occurs in the microaerobic ileum, and that *atcA*-mediated anaerobic androgen catabolism mainly occurs in the anaerobic caecum, where oxygen is often exhausted by gut microbes (Singhal and Shah, 2020).

The highest copy number of the *Thauera* 16S rRNA was observed in the caecum of the strain GDN1-administered mice, indicating that strain GDN1 mainly colonized in the mouse caecum. Moreover, in the anaerobic (denitrifying) culture spiked with the mouse feces, we noted that strain GDN1 considerably consumed testosterone within 3 days, indicating the high survival rate of strain GDN1 in the mouse gut. Our qPCR and RT-qPCR results indicated that strain GDN1 mainly colonized in the mouse caecum, which has been considered the main location where steroids such as cholesterol and bile salts (Adlercreutz et al., 1979; Cai and Chen, 2014; Lu et al., 2001) are reabsorbed through enterohepatic circulation. In animals, gut androgen content is typically quite low; to enable androgen catabolism, gut bacteria have to efficiently access, import, and accumulate this substrate from the environment. Steroid-catabolic bacteria often exhibit high cell surface hydrophobicity and high substrate uptake efficiency (Lin et al., 2015). Strain GDN1 may interrupt enterohepatic circulation through the adhesion, uptake, as well as aerobic and anaerobic catabolism of gut androgens (mainly testosterone), leading to low androgen reabsorption into blood. Thus far, the information on the substrate uptake mechanisms and corresponding transporters of androgen-catabolic bacteria remains unclear.

The strain GDN1 administration caused a minor increase in the relative abundance of *Thauera* (< 0.1%) in the gut microbiota. However, it profoundly affected the host physiology and gut bacterial community structure. Our data indicate that strain GDN1-mediated changes in bacterial community structure are sex-dependent. In the strain GDN1-administered mice, the decreased abundance of Akkermansiaceae, Bifidobacteriaceae, and Enterobacteriaceae populations was observed. These gut bacteria have been reported to associate with circulating testosterone levels in mice and humans (Mayneris-Perxachs et al., 2020). Moreover, members of Akkermansiaceae, Bifidobacteriaceae, and Enterobacteriaceae appear to play a role in prostate cancer progression (Doden et al., 2019; Huang et al., 2021). Our data thus indicate that strain GDN1 can shape the gut bacteria community through regulating androgen metabolism in the animal gut.

## Conclusions

The results of microbial community structure analysis suggest that androgen metabolism in mouse gut is achieved through synergistic microbial networks. In this study, we discovered the occurrence of androgen catabolism in the animal gut as well as the significant effect of this microbial activity on the regulation of host circulating androgens. Our biochemical and molecular data provided mechanistic insights into androgen-mediated gut microbe–host interactions along with the characteristic metabolites and functional genes for clinical diagnosis. This is important considering the future applications of androgen-catabolic bacteria as probiotics to reduce hyperandrogenism-associated host symptoms such as androgenetic alopecia, benign prostatic hypertrophy, and prostate cancer. Currently, androgen-deprivation therapy remains the mainstay of prostate cancer treatment. However, after an initial favorable response, patients often develop resistance to androgen-deprivation therapy, resulting in tumor progression, partially due to androgen production from steroidal precursors such as glucocorticoids and pregnenolone by some gut microbes (Doden et al., 2019; Huang et al., 2021; Pernigoni et al., 2021). Therefore, new therapeutic strategies are urgently required. In addition, the redox reactions of androgens occurring in the animal gut may alter circulating testosterone levels and thus lead to adverse effects on host physiology (Li et al., 2022). In the animal gut, which has a highly reducing environment, testosterone is often reduced to DHT, the most potent natural androgen (Colldén et al., 2019). Therefore, gut microbes responsible for androgen reduction are often considered unfavorable species (Li et al., 2022). Our data indicate the possible applicability of androgen-catabolic gut bacteria as potent probiotics in alternative therapy of hyperandrogenism.

### Significance

Denitrifying betaproteobacteria *Thauera* spp. are metabolically versatile and widely distributed in O_2_-limited ecosystems including the animal gut; however, only a subset of *Thauera* spp. harbors the androgen-catabolic capability. Here, we found that *Thauera* sp. strain GDN1 administration through oral gavage regulated mouse serum androgen levels, possibly because it blocked androgen recycling through enterohepatic circulation. We elucidated the mechanism underlying gut microbe–mediated androgen catabolism through biochemical, genetic, and metabolite profile analyses. Androgen catabolism in the mouse gut was noted to proceed through both the O_2_-dependent (mainly in the microaerobic ileum) and O_2_-independent (mainly in the anaerobic caecum) catabolic pathways. Our discovery portends a possibility of harnessing androgen-catabolic gut bacteria as functional probiotics to treat hyperandrogenism.

## Acknowledgements

This study was supported by the Ministry of Science and Technology of Taiwan (MOST 110-2311-B-001-033-MY3, 109-2314-B-002-125-MY3, 110-2811-B-002-562, and 110-2222-E-008-002) and Academia Sinica Career Development Award (AS-CDA-110-L13). We thank the Small Molecule Metabolomics core facility sponsored by the Institute of Plant and Microbial Biology (IPMB), Academia Sinica for UPLC–HRMS analysis.

## Authors contributions

T.-H.H., C.-H.C., M.-J.C., and Y.-R.C. designed research; T.-H.H., C.-H.C., Y.-L.C., G.-J.B.-M., T.-Y.W., P.-T.L., C.-W.L., Y.-L.L., and Y.-L.T. performed research; P.-H.W., T.-H.L., and C.-J.S. contributed new reagents/analytic tools; C.-H.C., Y.-L.C., M.-J.C. and Y.-R.C. analyzed data; and T.-H.H., P.-H.W. and Y.-R.C. wrote the paper

## Declaration of interests

The authors have no conflicts of interest to declare.

## Materials and Methods

### Chemicals

The [2,3,4C-^13^C]testosterone was purchased from Isosciences. Commercially available steroid standards [androstadienedione (ADD), androstenedione (AD), cholesterol, dihydrotestosterone (DHT), estradiol, 17α-ethinylestradiol, progesterone, and testosterone] were purchased from Sigma-Aldrich (St. Louis, MO, USA). The androgenic ring-cleaved metabolites 3,17-DHSA and 2,3-SAOA were produced as described elsewhere (Wang et al., 2013). Other chemicals used were of analytical grade and were purchased from Mallinckrodt Baker (Phillipsburg, USA), Merck Millipore (Burlington, USA), and Sigma-Aldrich unless specified otherwise.

### Detection of androgen metabolites and determination of androgenic activity in the denitrifying strain GDN1 culture

Denitrifying strain GDN1 (500 mL, initial OD_600 nm_ = 0.2) was incubated at 37°C and pH 6.0 with testosterone (1 mM) and nitrate (10 mM) in a chemically defined medium as described elsewhere (Shih et al., 2017). 17α-Ethinylestradiol (final concentration = 50 μM), which cannot be utilized by strain GDN1, was added to the bacterial culture to serve as an internal control. Cultural samples (5 mL each) were withdrawn every 8 h (0∼48 h). The pH of the cultural samples (1 mL) was adjusted to pH < 2 using 5 M HCl. The acid-treated cultures were extracted three times with the same volume of ethyl acetate to recover the residual testosterone and its derivatives from the aqueous phase. The testosterone-derived intermediates extracted from the cultural samples were identified and quantified using UPLC–HRMS. The protein and nitrate contents of strain GDN1 cultures were determined as described below. The remaining cultural samples (1 mL) were extracted with ethyl acetate three times. After the solvent was completely evaporated, the residue was re-dissolved in 10 μL of dimethyl sulfoxide (DMSO) to determine its androgenic activity (see below).

### Aerobic growth of strain GDN1 on testosterone

Strain GDN1 was aerobically grown in a phosphate-buffered shake-flask (300 ml in 1 L-Erlenmeyer flask) containing 1 mM testosterone. 17α-Ethinylestradiol (50 μM; indigestible by strain GDN1) was added as an internal control. Nitrate was omitted from the medium. In 1 L of distilled water, the medium contained the following: 0.29 g testosterone, 2.0 g NH_4_Cl, 0.5 g MgSO_4_•7H_2_O, and 0.1 g CaCl_2_•2H_2_O. After autoclaving, sterile 50 ml KH2PO_4_-K2HPO_4_ buffer solution (1 M, pH 6.0), vitamins (1 mL/L) (Pfennig, 1980), EDTA-chelated mixture of trace elements (1 mL/L) (Rabus and Widdel, 1995), and selenite and tungstate solution (1 mL/L) (Tschech and Pfennig, 1984) were added. The aerobic culture was incubated at 37°C in an orbital shaker (180 rpm). The cultural samples (5 mL) were withdrawn every 6 h (0∼30 h). The testosterone-derived intermediates extracted from the cultural samples were identified and quantified using UPLC–HRMS. The androgenic activity of the cultural samples was determined as described later.

### Measurement of protein content and nitrate concentration

Strain GDN1 cultural samples or cell suspension (0.2 mL) were centrifuged at 10,000 × *g* for 10 min. After centrifugation, the cell pellet was resuspended in 1 mL of reaction reagent (Pierce BCA protein assay kit; Thermo Scientific). The protein content was determined using a BCA protein assay according to manufacturer’s instructions with bovine serum albumin as the standard. The supernatant (0.1 mL) was diluted with 0.9 mL of double-distilled water, and the nitrate was determined using the cadmium reduction method according to manufacturer’s instructions (Nitrate Reagent Kit HI93728-01, Hanna Instruments).

### *lac*Z-based yeast androgen bioassay

The yeast androgen bioassay was conducted as described elsewhere (Wang et al., 2014). Briefly, the individual androgen standards or cell extracts were dissolved in DMSO, and the final concentration of DMSO in the assays (200 μL) was 1% (v/v). The resulting DMSO solutions (2 μL) were added to yeast cultures (198 μL, initial OD_600nm_=0.5) in a 96-well microtiter plate. The β-galactosidase activity was determined after 18 h incubation at 30°C. The yeast suspension (25 μL) was added to a Z buffer (225 μL) containing *o*-nitrophenol-β-D-galactopyranoside (ONPG, 2 mM), and the reaction mixtures were incubated at 37°C for 30 min. After the reactions were stopped by adding 100 μL of 1 M sodium carbonate, the amount of yellow-colored nitrophenol product was measured at 420 nm on a microplate spectrophotometer.

### Analytical chemical methods

#### (a) Thin-layer Chromatography (TLC)

The steroid standards and products were separated on a silica gel-coated aluminum TLC plate (Silica gel 60 F_254_: thickness, 0.2 mm; 20 × 20 cm; Merck) using dichloromethane:ethyl acetate:ethanol (14:4:0.05, v/v/v) as the developing phase. The steroids were visualized under UV light at 254 nm or by spraying the TLC plates with 30% (v/v) H_2_SO_4_, followed by an incubation for 1 min at 100°C (in an oven).

#### (b) High-Performance Liquid Chromatography (HPLC)

A reversed-phase Hitachi HPLC system equipped with an analytical RP-C18 column [Luna 18(2), 5 μm, 150 × 4.6 mm; Phenomenex] was used for separating steroid metabolites in this study. The separation was achieved isocratically using a mobile phase of 45% methanol (v/v) at 35°C at a flow rate of 0.5 mL/min. The steroid metabolites were detected using a photodiode array detector (200–450 nm). In some studies, HPLC was also used for quantifying steroids extracted from the strain GDN1 cultures. The quantity of major androgens (AD, ADD, DHT, and testosterone) was determined using a standard curve generated from individual steroid standards. The *R*^2^ values for the standard curves were >0.98. The presented data are the average values of three experimental measurements.

#### (c) Ultra-Performance Liquid Chromatography–High-Resolution Mass Spectrometry (UPLC– HRMS)

The androgen metabolites in mouse fecal (0.1 g) or serum (0.2 mL) samples were extracted using ethyl acetate for three times. Before the extraction, 0.1 ng of [2,3,4C-^13^C]testosterone (internal standard) was added to the samples. After the solvent was evaporated, the residue was re-dissolved in 10 μL of methanol. The androgen metabolites were identified by comparing their corresponding retention time and *m/z* values to the authentic standards. The androgen metabolites were quantified using the standard curves established using the authentic standards (linear range: 0.05–20 ng/mL; 2-fold serial dilution).

Androgen metabolites were analyzed using UPLC**–**HRMS on a UPLC system coupled to either an Electric Spray Ionization**–**Mass Spectrometry (ESI**–**MS) system or an Atmosphere Pressure Chemical Ionization**–**Mass Spectrometry (APCI**–**MS) system. Androgen metabolites were firstly separated using a reversed-phase C18 column (Acquity UPLC^®^ BEH C18; 1.7 μm; 100 × 2.1 mm; Waters) at a flow rate of 0.4 mL/min at 35°C (oven temperature). The mobile phase comprised a mixture of two solutions: solution A [0.1% formic acid (v/v) in 2% acetonitrile (v/v)] and solution B [0.1% formic acid (v/v) in methanol]. Separation was achieved using a gradient of solvent B from 10% to 99% in 8 min. ESI**–**HRMS analysis was performed using a Thermo Fisher Scientific^TM^ Orbitrap Elite^TM^ Hybrid Ion Trap-Orbitrap Mass Spectrometer (Waltham, MA, USA). Mass spectrometric data in positive ionization mode were collected. The source voltage was set at 3.2 kV; the capillary and source heater temperatures were 360°C and 350°C, respectively; the sheath, auxiliary, and sweep gas flow rates were 30, 15 and 2 arb units, respectively. APCI–HRMS analysis was performed using a Thermo Fisher Scientific^TM^ Orbitrap Elite^TM^ Hybrid Ion Trap-Orbitrap Mass Spectrometer (Waltham, MA, USA) equipped with a standard APCI source. Mass spectrometric data in positive ionization mode (parent scan range: 50–600 *m/z*) were collected. The capillary and APCI vaporizer temperatures were 120°C and 400°C, respectively; the sheath, auxiliary, and sweep gas flow rates were 40, 5 and 2 arbitrary units, respectively. The source voltage was 6 kV and the current was 15 μA. The elemental composition of individual adduct ions was predicted using Xcalibur™ Software (Thermo Fisher Scientific).

### General molecular biological methods

The strain GDN1 genomic DNA was extracted using the Presto^TM^ Mini gDNA Bacteria Kit (Geneaid, New Taipei City, Taiwan). PCR mixtures (50 μL) contained nuclease-free H_2_O, 2 × PCR master mix (Invitrogen™ Platinum™ Hot Start PCR 2X Master Mix, Thermo Fisher Scientific, Waltham, MA, USA), forward and reverse primers (200 nM each), and template DNA (10-30 ng). The PCR products were verified using standard TAE-agarose gel (1.5%) electrophoresis with the SYBR^®^ Green I nucleic acid gel stain (Invitrogen Thermo Fisher Scientific, Waltham, MA, USA), and the PCR products were purified using the GenepHlow Gel/PCR Kit (Geneaid, New Taipei City, Taiwan). The TA cloning was performed with T&A™ Cloning Vector Kit (YEASTERN BIOTECH, New Taipei City, Taiwan).

### PacBio sequencing of the strain GDN1 genome

For library preparation and PacBio sequencing, approximately 1 μg of strain GDN1 genomic DNA was sheared by Covaris g-TUBE (Covaris, Woburn, MA, USA) and purified via AMPure PB beads (PacBio, Menlo Park, CA, USA). The sheared and purified DNA fragments were used as templates to prepare the SMRTbell library through SMRTbell template prep kit 1.0 (PacBio, Menlo Park, CA, USA), according to the manufacturer’s instructions. After adaptor-ligation to the inserts, the inserts with suitable size for sequencing were selected with the BluePippin system. The SMRT sequencing was performed on SMRT 1M Cell v3 (PacBio, Menlo Park, CA, USA) with chemistry version 3.0 on PacBio Sequel sequencer (Genomics BioSci & Tech Co). A primary filtering analysis was achieved on the Sequel System, and the secondary analysis was completed using the SMRT analysis pipeline version 8.0. For the genome assembly, the filtered subreads after SMRT Link v8.0 were assembled by a long-read assembly algorithm Flye v2.7 (Kolmogorov et al., 2019). The SSPACE-LongRead v1.1 (Boetzer and Pirovano, 2014) was applied for draft genome scaffolding and PBJelly v15.8.24 was used for gap closure (Boetzer and Pirovano, 2014; English et al., 2012). Subsequently, genome polishing was conducted with Arrow v2.3.3 software (PacBio, Menlo Park, CA, USA). A further assembly of the circularizing genome was conducted using Circlator v1.5.5 (Hunt et al., 2015). Finally, the quality of the assembled genome was evaluated by QUAST v4.5 (Gurevich et al., 2013). After the *de novo* genome assembly, the annotations of genomic bacterial features were achieved with Prokka v1.13 (Seemann, 2014).

### RNA extraction and comparative transcriptomic analysis

The transcriptomes were extracted from the strain GDN1 cells grown aerobically or anaerobically with acetate (10 mM) or testosterone (2 mM) as the sole carbon source. The transcriptomes were extracted using the Direct-zol RNA MiniPrep Kit (Zymo Research, Irvine, CA, USA), and were further purified using Turbo DNA-free Kit (Thermo Fisher Scientific, Waltham, MA, USA) to eliminate DNA. rRNA was removed from the transcriptome samples using the Ribo-Zero rRNA Removal Kit (Epicentre Biotechnologies; Madison, WI, USA). The quality of the resulting RNA library was assessed on a Bioanalyzer 2100 system (Agilent Technologies, CA, USA) by using the RNA Nano 6000 Assay Kit. Only the samples with an integrity value in the range between 8 to 10 were selected for further transcriptomic analysis. First-strand cDNA was synthesized using the purified transcriptomes as the templates. Second-strand cDNA was synthesized in a reaction mixture containing Second-Strand Synthesis Kit (New England Biolabs; Ipswich, MA, USA). cDNA was purified using the Qiagen purification kit, followed by end repair using NEBNext^®^ End Repair Module (New England Biolabs). Antisense strand DNA was digested in the samples using the Uracil-Specific Excision Reagent Enzyme Kit (New England Biolabs), followed by a PCR to amplify the cDNA. The constructed sequencing libraries were sequenced as paired-end reads (with 150–200 bp read length) on the Illumina NovaSeq 6000 system (Illumina; San Diego, CA). Raw reads were first processed through in-house scripts (Novogen, Singapore), and the adapters were removed. In addition, reads with uncertain nucleotides (N) more than 10% or reads with low-quality nucleotides (base quality < 5; constituting >50% of the reads) were discarded. More than 98.2% of qualified read pairs were kept for mapping against strain GDN1 genome using Bowtie2 (version 2.2.3). The mismatch parameter is set to two, and other parameters are set to default. Finally, more than 96% of the total reads were aligned to the strain GDN1 genome (**Table S1**). The mapping results were quantified using the python script *htseq-count* (https://htseq.readthedocs.io/en/master/) and gene expression levels were estimated using fragments per kilobase of transcript per million mapped reads (FPKM).

### Preparation of strain GDN1 cell suspension for mice administration

Strain GDN1 was aerobically grown in tryptone soy broth (700 mL), and the bacterial cultures were incubated at 37°C in an orbital shaker (180 rpm). The bacterial cells were harvested at log phase (OD_600 nm_ = 0.5∼0.7) through centrifugation, and the cell pellet was washed and resuspended in a denitrifying mineral medium (500 mL) containing 0.2 mM testosterone (the inducer of androgen catabolic genes). The denitrifying strain GDN1 cultures were incubated at 37°C with shaking. The strain GDN1 cultures were sampled (1 mL each) twice per day, and the residual testosterone was determined using HPLC. When testosterone was exhausted (approximately 3 days), the strain GDN1 cells were washed twice using basal mineral medium (composed of 4.0 g NH_4_Cl, 4 g MgSO_4_•7H_2_O, 0.8 g CaCl_2_•2H_2_O, 4.2 g NaHCO_3_ and 1.7 g KH_2_PO_4_ per liter of distilled water), and then resuspended in the same mineral medium (adjusted to OD_600 nm_ = 2). The colony-formation-units (CFUs) of the resulting cell suspension were determined by counting the numbers of strain GDN1 colonies grown on tryptone soy agar. This cell suspensions of strain GDN1 were stored at 4 °C (within 5 days) before use.

### GDN1 administration through oral gavage

C57BL/6J mice (aged 7-week-old) were obtained from the Animal Center of the Medical College (National Taiwan University, Taipei, Taiwan) and were kept in standard animal housing conditions with the health guidelines for the care and use of experimental animals. The experiments were approved by the local ethics committee (IACUC No. 20210326). After acclimatization for one week, female and male mice with similar body weight (16–18 g) were randomly assigned into two treatment groups [vehicle-administered mice (VEH) or strain GDN1-administered mice (+GDN1)]. For administration of stain GDN1, approximately 5 × 10^8^ CFUs (suspended in 200 uL of basal mineral medium) were fed into each mouse through oral gavage twice per week (**Fig. 4*A***). The same volume of sterilized basal mineral medium was administered to the vehicle mice. During the period of continuous administration, the drinking water for mice was supplemented with sodium nitrate (2mM). Fresh mouse feces were collected daily, and the body weight was measured once per week. Mice were sacrificed on day 25 (anesthetized with 3% isoflurane) and the mouse blood was collected through cardiac puncture. The mouse gastrointestinal (GI) tracts, including the ileum, caecum, and colon, were dissected to obtain the GI contents (for bacterial DNA and RNA extraction). All the mouse samples, including serum, feces, and GI contents, were stored at -80℃ before use.

### Determination of serum testosterone and estradiol levels using ELISA

The serum testosterone level of mice was determined using a competitive binding Testosterone Parameter Assay Kit (R&D Systems, Minneapolis, MN) according to the manufacturer’s instructions with testosterone as the standard. The minimum detectable dose of testosterone for the ELISA kit was approximately 0.03 ng/mL. The following compounds were tested for their cross-reactivity (testosterone cross-reactivity was set as 100%): DHT (2.6%), AD (< 0.1%), 17β-estradiol (< 0.1%), and progesterone (< 0.1%). On the other hand, the serum estradiol level of mice was determined using a competitive binding Estradiol Parameter Assay Kit (R&D Systems, Minneapolis, MN) according to the manufacturer’s instructions with 17β-estradiol as the standard. The minimum detectable dose of estradiol was approximately 5 pg/mL. The following compounds were tested for cross-reactivity (17β-estradiol cross-reactivity was set as 100%): estrone (0.26%), estriol (0.86%), 17α-ethinylestradiol (< 0.1%), and progesterone (< 0.1%).

### Extraction of bacterial DNA from mouse fecal samples and GI content

Bacterial DNA in the mouse fecal samples and GI content (200 mg) was extracted using QIAamp^®^ PowerFecal^®^ Pro DNA kit (Qiagen, Hilden, Germany) according to the manufacturer’s instructions. The DNA concentration was determined using NanoDrop Spectrophotometer ND-1000 or Qubit^TM^ dsDNA Assay kit (Invitrogen Thermo Fisher Scientific, Waltham, MA, USA).

### Extraction of bacterial RNA from mouse GI content and reverse transcription of the RNA

Bacterial RNA from mouse GI contents (approximately 250 mg) was extracted using RNeasy^®^ PowerMicrobiome^®^ kit (Qiagen, Hilden, Germany) according to the manufacturer’s instructions. To confirm the complete DNA elimination, the resulting RNA samples were amplified using bacterial universal 16S rRNA primers 27F and 1942R as described elsewhere (Shih et al., 2017). The RNA concentration was determined using NanoDrop Spectrophotometer ND-1000 or Qubit^TM^ HS RNA Assay kit (Invitrogen Thermo Fisher Scientific, Waltham, MA, USA). The quality of the bacterial RNA samples was examined based on the ratio of 23S/16S rRNA separated through agarose-electrophoresis (Farnsworth et al., 2004). Bacterial RNA (1 ug) was reverse-transcribed using SuperScript^TM^ IV kit (Invitrogen Thermo Fisher Scientific, Waltham, MA, USA) according to the manufacturer’s instructions. The resulting cDNA was used for the quantitative PCR (qPCR).

### Real-time qPCR

The bacterial DNA or cDNA was quantified through qPCR by using the *Power*^®^ SYBR Green PCR Master Mix (Applied Biosystem, Thermo Fisher Scientific, Waltham, MA, USA) on the QuantStudio 5 platform (Applied Biosystem, Thermo Fisher Scientific, Waltham, MA, USA) according to the manufacturer’s instructions. Primers used for the qPCR are shown in **Table S2**. To determine the abundance of *Thauera* species in the mouse feces and GI contents, a real-time qPCR-based assay (Kleindienst et al., 2017) was performed; a calibration curve was obtained by using 10-fold serial dilution of plasmid DNA carrying a cloned strain GDN1 16S rRNA gene. To quantify the expression of androgen catabolic genes *atcA* and *kshA* in the mouse GI tract (ileum, caecum, and colon), the relative change of gene expression was calculated using the 2^-ΔΔCt^ method (Livak and Schmittgen, 2001) with the Ct value of bacterial universal 16S rRNA (see **Table S2** for nucleotide sequences) as the internal control. The expression of individual androgen catabolic genes in the VEH ileum was set as 1.

### Bacterial 16S rRNA sequencing and data processing

For bacterial 16S rRNA sequencing, the hypervariable (V3-V4) region was amplified using the primer set (16S-341F/805R, see **Table S2** for nucleotide sequences) according to the 16S Metagenomic Sequencing Library Preparation procedure (Illumina; San Diego, CA). The success of PCR amplification was confirmed using agarose electrophoresis (1.5%), and the PCR products with a size of approximately 500-bp were selected and purified using the AMPure XP beads. The resulting PCR products were ligated to the dual indices and Illumina sequencing adapters with Nextera XT Index Kit. The quality of indexed PCR products was examined on the Qubit 4.0 Fluorometer (Invitrogen Thermo Fisher Scientific, Waltham, MA, USA) and Qsep100TM Capillary electrophoresis system (Bioptic Inc. New Taipei City, Taiwan). The purified indexed PCR products were further processed according to the Illumina standard protocol, and sequenced on an Illumina MiSeq platform (paired-end 2 × 300 bp) at the BIOTOOLS Co., Ltd (New Taipei City, Taiwan).

The raw Illumina amplicon reads were quality-filtered and analyzed using mothur v1.46.1 according to mothur MiSeq standard operating procedure (Kozich et al., 2013). In brief, the forward and reverse Fastq files were merged; the resulting sequences with low quality (quality score < 30), merged sequences with a length < 400 bp or > 600bp, sequences containing nucleotide repeats (> 8), and primer sequences were all removed from the dataset. The sequences were further denoised using *pre.cluster* command and amplicon sequence variants (ASVs) were generated with two base-pair differences. Chimeric sequences were removed using VSEARCH algorithm (*chimera.vsearch*) within mothur. The ASVs were assigned into a taxonomic hierarchy based on the reference sequences from the SILVA database (version 138_1) using mothur (*classify.seqs*). Sequences classified as chloroplast, eukaryote, mitochondria, or unknown were removed from the dataset (*remove.lineage*). To normalize the dataset, each sample was rarefied to the minimum of 48558 for downstream analysis (*sub.sample*).

### Gut bacterial community structure analysis and the construction of microbial interaction networks

To evaluate beta-diversity, we used non-metric multidimensional scaling (NMDS) ordination approach based on Bray-Curtis dissimilarities to assess the composition of bacterial communities between the vehicle and strain GDN1 treatments. The NMDS was calculated using the *vegan* package in R studio. The significant changes in bacterial community structure were assessed using permutational analysis of variance (PERMANOVA) in R studio using the package *vegan*. DESeq2 was used to analyze the differential abundance of bacterial taxa (i.e., Phylum and Family levels) between the vehicle and strain GDN1 treatments based on Benjamini-Hochberg adjusted p-value to reduce false discovery rate (FDR) (Love et al., 2014). The top 5 highly abundant bacterial phyla were visualized as histograms, whereas the bacterial families that were significantly different between groups were visualized as heatmaps using ClustVis (Metsalu and Vilo, 2015). To simulate potential interactions between different bacterial genera, we generated SparCC networks with 1000 permutations based on centered log-ratio transform data and spearman correlations (Friedman and Alm, 2012) in R studio using the package *SpiecEasi* (Kurtz et al., 2015). SparCC values with *p*-value less than 0.05 were considered significant, and were selected to generate bacterial interaction networks in Gephi (Bastian et al., 2009). Node centralities such as degree, closeness, betweenness, and eigenvector were calculated in Gephi (Bastian et al., 2009). The bacterial genera with potential important roles in a network were selected based on their eigenvector centrality values. (Peschel et al., 2021). A high eigenvector centrality value indicates relatively high degree, closeness, and betweenness centrality values. The top 30 bacterial genera with the highest eigenvector centrality value were selected to visualize the network. Eigenvector centrality is indicated by node size and color: a larger round node with a darker red hue corresponds to a higher value of eigenvector centrality. Conversely, a smaller round node with a bright yellow hue corresponds to a lower value of eigenvector centrality. A red edge depicts a positive interaction, while a blue edge depicts a negative interaction.

## Supplementary Information

### Additional file 1

#### Supplemental Results SI Figures

**Fig. S1** Determination of relative gene expression of the strain GDN1 *atcA* (**A**) and *kshA* (**B**) using RT-qPCR.

**Fig. S2** The administration of mice with strain GDN1 through oral gavage for 25 days did not apparently change the host serum estradiol level of male mice.

**Fig. S3** Bacterial community structure analysis indicated the temporal changes in the relative abundance of *Thauera* in fecal samples.

**Fig. S4** KEGG (Kyoto Encyclopedia of Genes and Genomes) analysis revealed a complete set of genes participating in denitrification (**A**) and TCA cycle (**B**).

**Fig. S5** The microbial interaction networks identified in the female (**A**) and male (**B**) mice.

**Fig. S6** The predicted interactions between *Thauera* and other gut microbes in the female (**A**) and male (**B**) mice.

#### SI Tables

**Table S1** Quality control of the RNA sequencing and the mapping status of clean reads.

**Table S2** Oligonucleotide primers used in this study.

**Table S3** Selection of housekeeping genes used as reference genes (black spots) in the global gene expression profiles of strain GDN1 (Figure 2C).

### Appendices

**Appendix S1.** Nucleotide sequence of the 16S rRNA gene (CKCBHOJB _02702) of strain GDN1.

**Appendix S2.** Nucleotide sequence of the *atcA* gene (CKCBHOJB _02411) of strain GDN1.

**Appendix S3.** Nucleotide sequence of the *kshA* gene (CKCBHOJB _02279) of strain GDN1.

**Appendix S4.** Nucleotide sequence of the inherent *atcA*-like DNA fragment extracted from the vehicle mouse caecum.

#### Additional file 2

**Dataset S1 (separate file).** Genome annotation of strain GDN1 and transcriptomic analysis (RNA-Seq) of bacterial cells grown with testosterone or acetate under both aerobic and anaerobic conditions. **Dataset S2 (separate file).** Eigenvector centrality values of individual bacterial genera in the gut microbiota of vehicle-administered and strain GDN1-administered female and male mice.

## Notes

### Competing Interest Statement

The authors have declared no competing interest.

